# Persistent CD4^+^ T cell functional deficits during recovery from prolonged symptomatic SARS-CoV-2 infection

**DOI:** 10.64898/2025.12.03.692161

**Authors:** Jadith Ziegler, Christina Lawrence, Sean Turner, Nathan Pezant, Nancy Redinger, Carla Guthridge, Kenneth Smith, Jingrong Chen, Joel M. Guthridge, Judith A. James, Susannah Rankin, Linda F. Thompson, A. Darise Farris, Susan Kovats

**Affiliations:** Arthritis & Clinical Immunology Program, Oklahoma Medical Research Foundation, 825 NE 13th Street, Oklahoma City, OK, USA; Center for Biomedical Data Sciences, Oklahoma Medical Research Foundation, 825 NE 13th Street, Oklahoma City, OK, USA; Cell Cycle and Cancer Biology Program, Oklahoma Medical Research Foundation, 825 NE 13th Street, Oklahoma City, OK, USA; Department of Microbiology & Immunology, University of Oklahoma Health Campus, 1100 N Lindsay Ave, Oklahoma City, OK, USA; Department of Pathology, University of Oklahoma Health Campus, 1100 N Lindsay Ave, Oklahoma City, OK, USA; Department of Cell Biology, University of Oklahoma Health Campus, 1100 N Lindsay Ave, Oklahoma City, OK, USA

**Keywords:** SARS-CoV-2, CD4 T cells, T cell dysfunction

## Abstract

Symptoms of acute SARS-CoV-2 infection often resolve quickly but are sometimes associated with persistent immune dysfunction. The factors that predispose individuals to compromised immune function have not been well defined. We investigated CD4^+^ T cell phenotype and function in a small cohort of individuals who recovered from mild to moderate SARS-CoV-2 infection without hospitalization and were divided into short or prolonged symptom duration groups. Five individuals with prolonged symptom duration showed marked downregulation of CD4 on CD3^+^CD8^−^ T cells (CD4^low^ group) and a poor response to TCR stimulation with the superantigen Staphylococcal enterotoxin B (SEB), as shown by weak upregulation of the activation markers CD134 and CD69. CD4 surface intensities recovered to normal levels in four of these individuals within 3-12 months. Selected cytokines (IL-1RA, IL-7, and VEGF) were elevated in individuals with low CD4, but plasma levels of anti-S1 IgG did not correlate with CD4 defects. Bulk RNA sequencing of unstimulated and SEB-treated CD3^+^CD8^−^ T cells revealed a >50% reduction in the number of differentially expressed genes in the CD4^low^ group compared to the same individuals after CD4 levels were recovered and a healthy control group. Upstream regulator analysis of differentially expressed genes in unstimulated CD4^low^ cells suggested a response to IFN, while SEB-stimulated CD4^low^ cells showed reduced functionality of IL-2, CD28, and SATB1 regulated pathways. In summary, prolonged symptomatic recovery from SARS-CoV-2 infection was associated with a global CD4^+^ T cell response defect, defined by low surface CD4 expression, evidence of IFN signaling, and defective T cell activation.

## 1. Introduction

SARS-CoV-2, the causative agent of the COVID-19 pandemic, primarily targets the respiratory tract but can result in varied clinical manifestations, ranging from asymptomatic or mild infections to severe cases linked to acute respiratory distress syndrome and death [1]. More than 80% of all individuals with COVID-19 experience mild to moderate symptoms, are managed as outpatients or through self-isolation at home, and are less studied than those with severe disease [2–4]. A subset of non-hospitalized individuals with acute COVID-19 shows prolonged symptoms and delayed recovery, which may be secondary to immune dysfunction induced by SARS-CoV-2. New studies have begun to link particular immunological features of the acute innate and adaptive responses to SARS-CoV-2 infection to the development of durable sequelae of COVID-19 [5–10]. Combined with clinical signs, identification of the spectrum of lymphocyte phenotypes induced upon acute SARS-CoV-2 infection will help determine whether specific types of lymphocyte responses contribute to prolonged immune dysfunction that may impact susceptibility to new infections or vaccine responses. Collecting this information will help inform treatment options, an important goal since the long-term consequences of COVID-19 are a significant burden to US and global public health [11].

The immune response to natural SARS-CoV-2 infection engages both the innate and adaptive immune systems. The innate antiviral response includes the release of chemokines, interferons, and cytokines by activated myeloid cells and lymphocytes. Viral proteins can dampen immunity by inhibiting type I interferon (IFN-I) production [12], and in severe cases, the lack of early viral clearance due to reduced IFN-I leads to excessive cytokine release, potentially causing damage and failure in various organ systems [13]. The adaptive immune response to SARS-CoV-2 that occurs early after symptom onset includes the activation of B cells and antigen-specific CD8^+^ and CD4^+^ T cells, which promote protective immunity by contributing to immediate viral clearance, antiviral antibody production, and long-term cellular immunity [14]. In mild COVID-19 cases, SARS-CoV-2-specific CD4^+^ T cells produce IFNγ and show phenotypic features of robust helper function and longevity, consistent with dominant T helper cell (T_H_1) and T follicular helper cell (T_FH_) responses [15]. In contrast, severe infections with poor outcomes are associated with a paucity of SARS-CoV-2-specific CD4^+^ T_H_1 and T_FH_ cells and a higher number of SARS-Cov-2-specific T regulatory cells [16]. The persistence of SARS-CoV-2-specific T cells for over two months post-infection [15, 17] and up to 22 months post-symptom onset [18] indicates the potential for enduring T cell-mediated immunity, although further research is necessary to fully understand the role of these memory T cells in durable protection against mutating viruses and the role of potential immune abnormalities. Indeed, long-term sequelae of COVID-19 have been correlated with persistent tertiary lymphoid organs in the lung, enduring immune activation, lymphocyte exhaustion and senescence, and dysregulated dynamics of viral antigen-specific B and T cells [8, 19, 20].

The chronic lymphocyte activation observed following acute COVID-19 may be attributed to several factors related to SARS-CoV-2 persistence and its unique structural features. The virus, its RNA, or antigens may remain in tissue reservoirs, continually stimulating the immune system. [21]. Additionally, the SARS-CoV-2 spike 1 (S1) glycoprotein harbors a unique superantigen-like motif resembling the superantigen Staphylococcal enterotoxin B (SEB) [22, 23]. Superantigens bind to specific TCR V beta regions, triggering CD4^+^ T cell activation without the need for a cognate MHC class II/peptide complex [24]. Superantigen binding allows for potent and widespread T cell activation, potentially contributing to the prolonged immune response seen in some individuals with COVID-19. The consequences of this chronic stimulation are multifaceted and can significantly impact T cell function. Computational analyses showed that the SEB-like motif in S1 binds to TCRs and MHCII proteins with high affinity [23], suggesting that the released S1 protein could lead to chronic CD4 T cell stimulation.

Chronic stimulation has been associated with a reduction of CD4 on T cells and altered magnitude and pathways of TCR signaling [25, 26]. This CD4 reduction is particularly significant as CD4 plays a crucial role in stabilizing the interaction between the T cell receptor and MHCII on antigen-presenting cells, thereby amplifying TCR signaling through the CD4-associated kinase LCK [27]. Furthermore, like SEB, the superantigen-like properties of the S1 protein may induce LCK-independent TCR signaling [28, 29]. These alterations in T cell signaling and function, combined with the persistent presence of viral antigens or the ongoing effects of interferons and cytokines triggered by residual viral RNA, may contribute to T cell hypo-responsiveness or exhaustion during the recovery phase from COVID-19.

In this study, we investigated the CD4^+^ T cell phenotype and function in individuals who had recovered from mild to moderate SARS-CoV-2 infection without hospitalization. We conducted a comprehensive analysis using blood samples from a small cohort of such individuals, several of whom were followed up longitudinally. Participants were divided into two distinct groups based on the duration of their symptoms: a “Prolonged” symptom group and a “Short” symptom group. This stratification enabled us to compare the immunological profiles of these two cohorts with each other and with those of a control group of healthy individuals who had not been exposed to SARS-CoV-2. The results revealed a persistent global defect in T cell CD4 expression and dysfunctional TCR signaling following prolonged symptomatic infection. The presence of the defect correlated with elevated proinflammatory cytokines such as VEGF, which is associated with COVID-19 severity [30], and evidence of T cell exposure to type I and II interferons.

## 2. Materials and Methods

### 2.1 Study design and participant cohort

This study was conducted in accordance with the principles of the Declaration of Helsinki. Ethical approval was obtained from the Oklahoma Medical Research Foundation (OMRF) Institutional Review Board (IRB#20-11). All participants provided written informed consent for sample collection and subsequent analysis. This study examined 17 initial samples from donors with prior SARS-CoV-2 infection, collected at OMRF between April 2020 and March 2021. The self-reported demographic composition of the study group comprised white participants, 41% of whom had Hispanic ethnicity, including both males and females aged 26–71 years (**Table 1, Table S1**). All participants had recovered from SARS-CoV-2 infection without hospitalization and were unvaccinated at the time of the initial sample collection. The participant samples were categorized into either a “Prolonged Symptom” group (n=9) or “Short Duration Symptom” group (n=8) based on symptom duration in days (median=30 vs. 5) and number of symptoms (median=9 vs. 3.5). Symptoms queried included fever, cough, loss of taste, fatigue, headache, chest pain/tightness, nausea/vomiting, chills, shortness of breath, loss of smell, body aches, dizziness, night sweats, and diarrhea. Peripheral blood from age-, race-, and sex-matched healthy control volunteers had been collected and banked at OMRF prior to 2019. Peripheral blood mononuclear cells (PBMCs) were isolated using Ficoll-Hypaque density gradient centrifugation and cryopreserved using standard methods.

**Table 1.** Study Participant Demographics. Self-reported demographics of 17 study participants and 12 healthy controls including both males and females aged 26-71. All study participants had recovered from SARS-CoV-2 infection and did not require hospitalization. They were categorized into either a “prolonged” symptom duration group or “short” duration group. Peripheral blood from healthy controls was banked at OMRF prior to 2019. “*ns”, not significant*.

| Parameters | Prolonged Duration Symptom Group (n=9) | Short Duration Symptom Group (n=8) | P-value | Healthy Controls (n=12) |
| --- | --- | --- | --- | --- |
| Age (mean± SD) | 51± 15.3 | 42.3±17.3 | ns<sup>a</sup> | 46.3±15.7* |
| Race (% White) | 9/9 (100%) | 8/8 (100%) | ns<sup>b</sup> | 10/12 (83%) |
| Ethnicity (% Hispanic) | 3/9 (33%) | 4/8 (50%) | ns<sup>b</sup> | 1/12 (9%) |
| Gender |  |  |  |  |
| *Female* | 6/9 (66%) | 4/8 (50%) | ns<sup>b</sup> | 8/12 (67%) |
| *Males* | 3/9 (34%) | 4/8 (50%) |  | 4/12 (33%) |
| Symptom Duration Days (median, range) | 30 (18-61) | 5 (4-7) | <0.0001<sup>c</sup> | NA |
| Number of symptoms (median, range) | 9 (3-11) | 3.5 (2-10) | 0.05<sup>b</sup> | NA |
| *cough* | 78% | 12.50% | 0.015<sup>b</sup> | NA |
| *shortness of breath* | 67% | 12.50% | 0.05<sup>b</sup> | NA |
| Sampling, Days post-recovery (median, range) | 29 (10-164) | 25 (9-174) | ns<sup>c</sup> | NA |

<sup>a</sup> Unpaired t-test, <sup>b</sup> Fisher's exact test, <sup>c</sup> Mann-Whitney U test
\*Mean ages for 10/12 HC donors. Age information was not available for 2/12 donors.

### 2.2 In vitro Superantigen (SEB) stimulation of thawed PBMC obtained from COVID-19 infected and control participants

Cryopreserved PBMCs from the donors were quickly thawed in a 37°C water bath. Cells were slowly added to pre-warmed AIM-V medium (Thermo Fisher Scientific) containing Benzonase nuclease (1:10,000 dilution; Sigma-Aldrich) to prevent cell clumping. Viable cells were counted using the trypan blue exclusion assay. PBMCs were resuspended at 6.7 × 10^6^ cells/mL in AIM-V medium containing 50 ng/ml FLT3 Ligand, and 150 μl were added to each well of a 96-well round-bottomed plate. After 24 h, Staphylococcal enterotoxin B (SEB; Fisher Scientific) was added to the appropriate wells at a final concentration of 1 μg/mL in 50 μl, while 50 μl of AIM-V medium was added to the “unstimulated” wells. The cells were then incubated for an additional 18 h at 37°C. Cells were harvested, washed with Dulbecco’s PBS, and stained with BV510 viability dye (BD Biosciences) for 15 min at room temperature. Fc receptors were blocked with Human BD Fc Block (BD Biosciences) for 10 min at room temperature prior to mAb staining.

### 2.3 T Cell Activation-induced marker assay

For the detection of CD4 and other markers on CD3^+^CD8^−^ T cells using flow cytometry, samples were stained with a cocktail of fluorochrome-conjugated antibodies for 30 min on ice. The antibody panel included: CD3-BB515 (BD Biosciences, clone UCHT1) CD4-BV711 (BD Biosciences, clone SK3), CD8-BV605 (BD Biosciences, clone SK1), OX40/CD134-APC (BD Biosciences, clone ACT35) and CD69-PerCP-Cy5.5 (Biolegend, clone FN50). Brilliant Stain Buffer Plus (BD Biosciences) was used to prevent fluorochrome interaction. After staining, the cells were washed and resuspended in FACS buffer (PBS + 5% FBS with no azide) for analysis. Compensation controls were prepared using single-stained cells and antibody capture beads. Data were acquired using LSR II or FACSAriaIIIu instruments (BD Biosciences).

### 2.4 Bulk Cell Sorting and Analysis

Unstimulated or SEB-stimulated CD3^+^CD8^−^ T cells were isolated by fluorescence-activated cell sorting (FACS) using the antibody panel described above and a FACSAriaIIIu cell sorter (BD Biosciences). Sorted cells were collected in FACS buffer for downstream analysis. Data were analyzed using FlowJo software (BD Biosciences). Gating strategies were employed to identify viable CD3^+^ T cells and further subsets of the population based on CD4, CD8, and activation markers.

### 2.5 Cytokine measurements

Plasma cytokine levels were measured using custom multiplex Luminex assays (R&D Systems) to quantify the levels of APRIL/TNFSF13, BMP2, CCL3/MIP1α, CXCL9/MIG, Fas Ligand/TNFSF6, IL1β/IL1F2, IL6, IL8/CXCL8, IL23, LIF, SCF/c-kit Ligand, TRAIL/TNFSF10, BAFF/BLyS/TNFSF13B, CCL2/MCP1, CCL4/MIP1β, CXCL10/IP10/CRG2, IFNγ, IL1Rα/IL1F3, IL7, IL10, IL33, PDGFββ, TNFα, VEGF, CCL7/MCP3/MARC, ICAM-1/CD54, IL5, β-NGF, Leptin/OB, GCSF, IL4, IL21, VCAM-1/CD106, CD40 Ligand/TNFSF5, IFNβ, IL13, E-Selectin/CD62E, CCL11/Eotaxin, IFNα, IL12 p70, Resistin, TNFRII/TNFRSF1B, Fas/TNFRSF6/CD95, IL2, IL17/IL17A, TNFRI/TNFRSF1A, CD25/IL2Rα, IL1α/IL1F1, IL15, Syndecan1/CD138, Oncostatin M/OSM, and TGFβ1. Data were analyzed using the Bio-Rad BioPlex200® array system (Bio-Rad Technologies) with a lower threshold of 50 beads per analyte/sample. The median fluorescence intensity (MFI) for each analyte was interpolated from 5-parameter logistic nonlinear regression standard curves. Analytes with levels below the detection limit were assigned values of 0.001 pg/ml.

### 2.6 SARS S1 antibody measurements

To determine the SARS-CoV-2 S1 antibody levels in plasma, high-binding ELISA plates (Costar 3369) were coated overnight at 4°C with 0.2 µg/well of SARS-CoV-2 S1 (Wuhan strain, expressed in-house from plasmid pCMV3-C-His [2019-nCoV Spike S1] (Sino Biological) in HEK293 cells).

Between each subsequent step, the plates were washed four times with PBS-0.05% Tween. After coating, the plates were blocked with 150 μl/well of 0.1% BSA in PBS for 1 h at room temperature, and plasma samples were loaded onto the plates. A fully human antibody against the SARS-CoV-2 spike protein, AM001414 (Amgen), was used starting at 5 μg/ml and serially diluted (4-fold dilutions) to generate a standard curve. The samples (1:100 and 1:400 dilutions) and standard curves were incubated on plates for two hours at room temperature and then washed. HRP-conjugated goat anti-human IgG (75 μl/well; Jackson ImmunoResearch) diluted 1:1000 was then added and incubated for 1 h at 37°C, followed by 100 μl/well of Super Aqua Blue ELISA substrate (Invitrogen). OD405 was measured using a plate reader. All samples were analyzed in at least three independent experiments on three different days, and the results were averaged.

### 2.7 Bulk RNA Sequencing

CD3^+^CD8^-^ T cells sorted from both SEB-stimulated and unstimulated conditions were assessed by bulk RNA Sequencing. RNA was isolated using the RNAeasy mini kit (Qiagen) according to the manufacturer’s instructions. RNA was quantified using a Fisher Qubit 4 Fluorometer (Thermo Fisher Scientific), and its quality was assessed using a High Sensitivity RNA ScreenTape (#5067-5579, Agilent Technologies) with a 4150 Tapestation analyzer (Agilent Technologies). Libraries were prepared using the xGen Broad-Range RNA kit (IDT) according to the manufacturer’s protocol. Briefly, mature mRNA was enriched from total RNA via pull-down with beads coated with oligo-dT homopolymers using the NEBNext Poly(A) mRNA Magnetic Isolation Kit (NEB). The mRNA molecules were then chemically fragmented, and the first strand of cDNA was generated using random primers that incorporated a truncated i5 adapter sequence. Following a bead-based cleanup, the 3’ end of the single-stranded cDNA was ligated to a truncated i7 adapter.

Libraries were then indexed using xGen Unique Dual Indexing primers (IDT). Final libraries for each sample were assayed on a Tapestation 4200 analyzer for appropriate size and quantity. These libraries were then pooled in equimolar amounts, as ascertained via fluorometric analyses. Final pools were quantified using qPCR on a QuantStudio6 instrument (ABI) with Illumina Library Quantification reagents (NEB). Sequencing of the libraries was performed on a NovaSeq 6000 (Illumina) with PE150 reads at the OMRF Clinical Genomics Core Facility. The RNA sequencing data were deposited in Mendeley: Kovats, Susan; Farris, A. Darise (2025), “CD4 T cell covid-19 Ziegler et al”, Mendeley Data, V1, reserved doi: 10.17632/8p2pzwbgw5.1.

### 2.8 Differential Expression Analysis

Differential expression (DE) analyses were conducted i) between unstimulated and SEB-stimulated conditions within each group (CD4^low^, CD4^rec^ and HC) separately and ii) between groups of subject-matched SEB-stimulated and unstimulated (no SEB) samples. Prior to analysis, the data quality was assessed using FASTQCv0.11.5. Low-quality reads were eliminated using Trimmomatic V.0.35. STAR v2.6.1c was used to align the FASTQ files to the human genome GRCh38. Furthermore, transcripts were excluded if either a) the count sum across all subjects was zero or b) the mean of the counts across all subjects was less than five, resulting in 19,826 transcripts remaining for the differential expression analysis. Analyses were performed utilizing the “results” function in DEseq2 package v1.36.0 in R v3.6.0, with the cooksCutoff option set to FALSE. Transcripts were considered differentially expressed if the results demonstrated a log_2_FC > 1.0 and an FDR-adjusted *P*-value <0.05.

### 2.9 Ingenuity Pathway Analysis (IPA)

Ingenuity Pathway Analysis software (IPA) (Qiagen) was used to identify the molecular pathways altered between the stimulated and unstimulated CD4^low^ and CD4 recovered (CD4^rec^) groups. Differentially expressed genes (DEGs) with log_2_FC ≥ ±0.58 (fold change 1.5) and FDR < 0.05 (padj <0.1) were analyzed using IPA’s core analysis function and the Ingenuity Knowledge Base. Significantly enriched pathways were identified using Fisher’s exact test (*P* < 0.05). Upstream regulators were predicted using IPA’s Upstream Regulator Analysis tool, with activation states determined by z-scores, with scores from -2 to +2 considered significant). Potential driver genes were identified using IPA Causal Network Analysis, considering the observed expression changes and known regulatory relationships. Significant predicted drivers had *P*-values < 0.05 and z-scores ranging from -2 to +2. All analyses used IPA version (01-22-01). The results were visualized using IPA’s Path Designer tool.

### 2.10 Statistical Analysis

Statistical analyses and visualizations were performed using Prism software version 9 (GraphPad). Differences in baseline characteristics between participants with persistent and short-duration disease were assessed using Fisher’s Exact Test for categorical variables and the Student’s t-test (paired or unpaired) or 1-way ANOVA for continuous variables. In figures 1 and 2, significance is denoted for the 1-way ANOVA as \**P* < 0.05, \*\**P* < 0.01, \*\*\**P* < 0.001, and for the paired t-test as ^#^*P* < 0.05.

**Figure 1.**
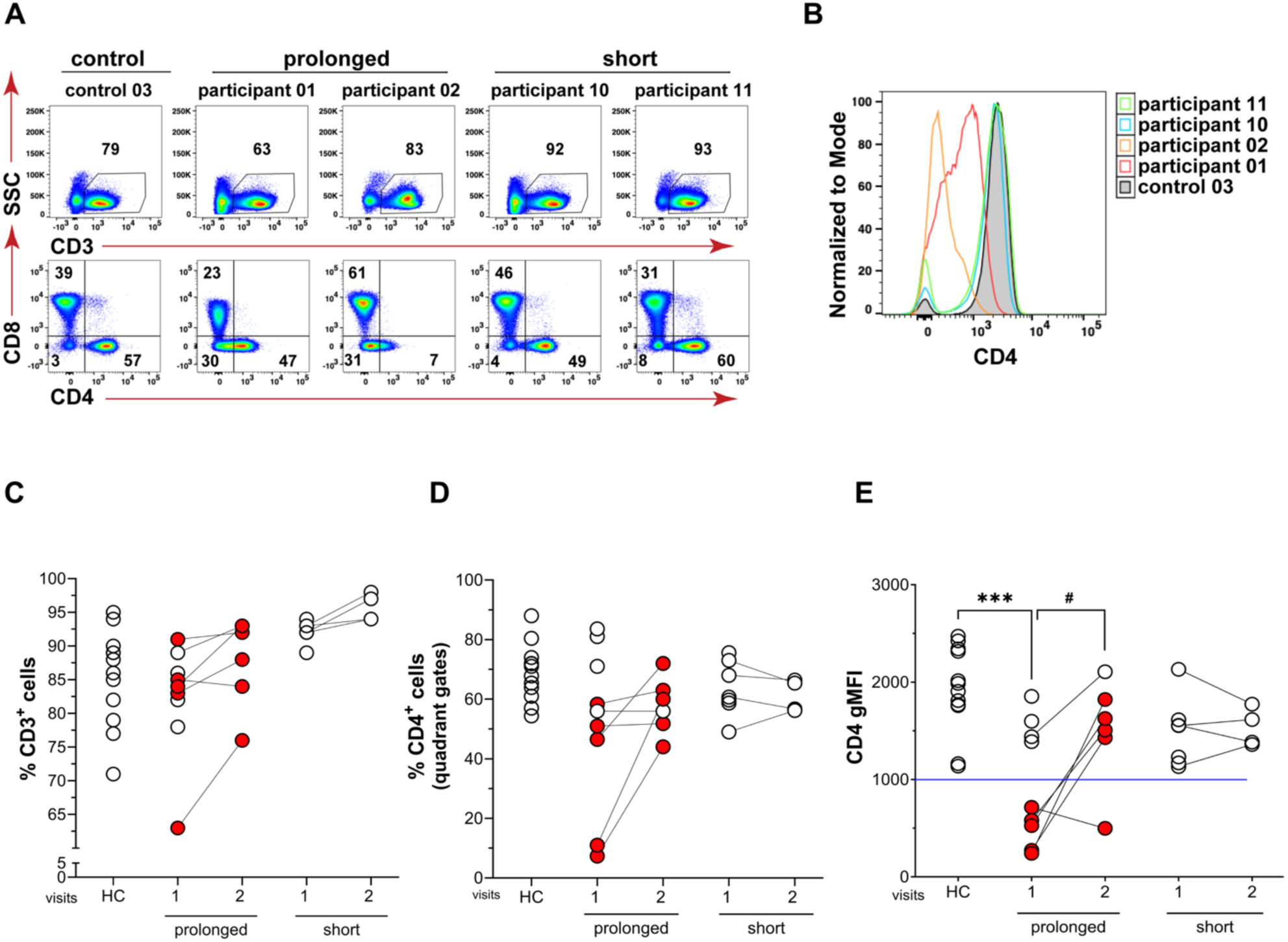
Downregulation of CD4 on CD3^+^ T cells in a subset of individuals with prolonged symptoms after SARS-CoV-2 infection. **(A)** Identification of CD3^+^CD4^+^ and CD3^+^CD8^+^ T cells within the PBMCs of study participants using flow cytometry. **(B)** Representative histograms showing CD4 levels on CD3^+^CD8^−^ cells. **(C)** % CD3^+^ T cells in live PBMCs of healthy controls (HC) and individuals recovered from prolonged or short duration of COVID-19 from two sequential blood draws (1, 2). **(D)** % CD3^+^CD8^−^ T cells based on quadrant gating in panel A. **(E)** CD4 gMFI (geometric mean fluorescence intensity) on CD3^+^CD8^−^ T cells defined in the quadrant gating scheme. The blue line indicates 2 SD from the mean of the control values. For all graphs, red symbols indicate the 5participants with a low CD4 gMFI in visit 1. Paired t-tests were conducted between blood draws 1 and 2 for each prolonged and short symptom group (*^#^, P < 0.05*). A one-way ANOVA was conducted with Tukey’s post-test to compare the HC group to each sample group and blood draw for each graph (*\*\*\*, P < 0.001*).

## 3. Results

### 3.1 Overview of Participant Cohorts

CD4^+^ T cell responses were examined in 17 individuals (males and females, ages 26-71) who had recovered from moderate SARS-CoV-2 infection between April 2020 and March 2021 without hospitalization **(Table 1, Table S1)**. Participants were grouped into “prolonged” (n=9) and “short” (n=8) cohorts based on symptom duration [median and range of 30 (18-61) days vs. 5 (4-7) days, respectively; *P*<0.0001], as well as the number of symptoms, such as coughing and shortness of breath [median and range of 9 (3-11) vs. 3.5 (2-10) symptoms, respectively; *P*=0.05]. Seven participants were studied at one time point, nine were studied at two time points, and one was studied at four time points. The time between blood draws ranged from 63 to 178 days. Three participants (01, 03, and 04) received their first COVID-19 vaccination before the second visit.

Participants’ results were compared to those of healthy controls (n=11) using blood collected prior to 2019 to ensure no SARS-CoV-2 exposure **(Table 1)**.

### 3.2 A subset of participants with prolonged symptoms after COVID-19 harbored CD3^+^CD8^−^ T cells bearing low levels of surface CD4

Flow cytometry was used to quantify the fraction of CD3^+^CD4^+^ and CD3^+^CD8^+^ T cells within the PBMCs of individuals who had recently recovered from COVID-19 in 2020 **(Fig. 1A, B)**. The percentage of CD3^+^ T cells was similar in both our prolonged and short patient groups and our control group **(Fig. 1A, C)**. However, in their initial blood draw, some individuals in the prolonged symptom group had markedly reduced percentages of CD3^+^CD4^+^ T cells **(Fig. 1A, D)** and a significant downregulation of CD4 compared to the control group, scored as geometric mean fluorescence intensity (gMFI), on CD3^+^CD8^−^ T cells **(Fig. 1E)**. Five participants (red symbols) with low CD4 levels (mean gMFI ± SD = 467 ± 204 vs. 1925 ± 443 for the control subjects; *P*<0.0001) in the prolonged-symptom group were studied after a second blood draw. In four out of five of these participants, T cells showed increased CD4 expression with the CD4 gMFI increasing from 406 ± 173 to 1596 ± 171 (*P=*0.0033). Notably, one participant did not show fully recovered CD4 expression over four visits **(Fig. S1A)**. In contrast, CD4 expression in the short-symptom-duration group remained unchanged between blood draws **(Fig. 1E)**.

### 3.3 T Cells with low CD4 Expression were hypo-responsive to TCR Stimulation

To determine whether T cells showing decreased surface CD4 had altered functional capacity, we stimulated PBMCs of the participants and healthy controls with SEB, a potent T cell superantigen that activates a large proportion of T cells via the TCR-Vβ chain and CD28 [24]. Using flow cytometry, we assessed T cell activation by measuring the percentages of cells expressing the early activation marker CD69 and the late activation marker CD134 **(Fig. 2A)** [31, 32]. Individuals harboring CD4^low^ T cells (indicated by the red symbol) during their first visit exhibited a markedly poor response to TCR stimulation, with significantly reduced percentages of CD134^+^CD69^+^ CD3^+^CD8^−^ T cells compared to the healthy control group (mean ± standard deviation: 12.7 ± 9.1 vs. 28.1 ± 6.9, *P=*0.0021) and compared to those with normal CD4 levels within the same prolonged symptom group (12.7 ± 9.1 vs. 28.7 ± 10.5, *P=*0.044) **(Fig. 2B)**. The percentages of CD134^+^CD69^+^ CD3^+^CD8^−^ T cells in the short duration group were similar to those in the healthy controls (28.1 ± 6.9 vs. 28.0 ± 4.2, *P=*0.97) **(Fig. 2B)**. Although we did not observe any statistically significant differences in T cell activation between the prolonged and short groups taken as a whole over their sequential visits, four of the five individuals with initial low CD4 expression showed an increase in their percentages of activated T cells after SEB stimulation on their second visit (9.17 ± 5.2 vs. 29.3 ±11.8, *P=*0.081) **(Fig. 2C)**, The percentages of cells expressing activation markers after SEB exposure strongly correlated with CD4 expression levels across all participant samples (Pearson’s test r = 0.74, *P*<0.0001) indicating that reduced CD4 expression is an effective biomarker for T cell dysfunction in this setting **(Fig. 2D)**. The recovery of CD4 expression was accompanied by the normalization of functional responses to SEB stimulation, with restoration of CD69 and CD134 upregulation (**Fig. 2C**). One individual (participant 04) exhibited a particularly delayed recovery, with persistent CD4 downregulation and functional deficit at 8 months post-infection, only beginning to recover at 11 months **(Fig. S1)**.

**Figure 2.**
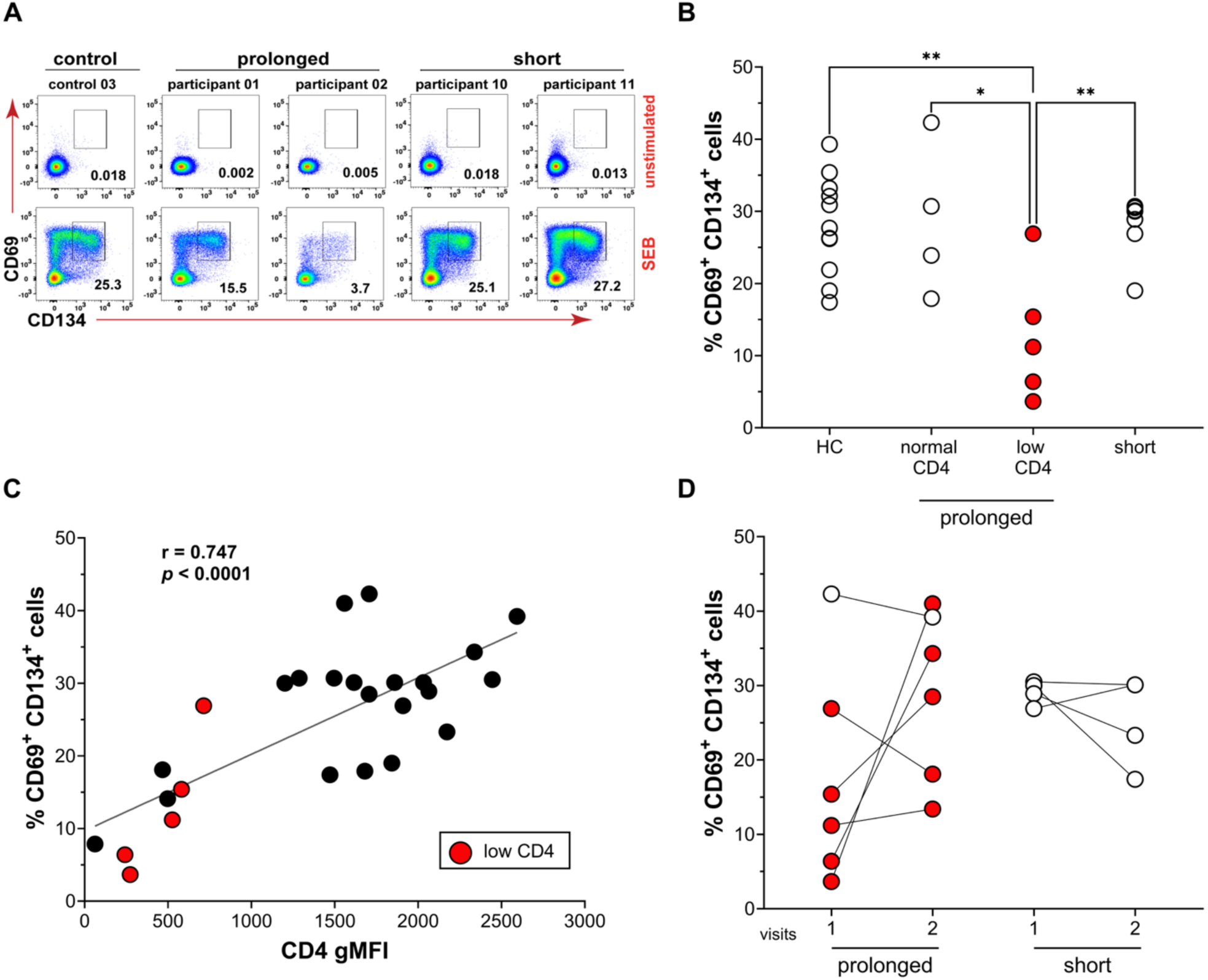
Lower CD4 expression levels in individuals with prolonged symptoms correlates with reduced T cell activation after SEB stimulation. **(A)** Representative CD69 vs. CD134 dot plots of CD3^+^CD8^−^ T cells from healthy controls and study participants before and after SEB stimulation. **(B)** Percentages of CD69^+^CD134^+^ activated CD3^+^CD8^−^ T cells after division of the prolonged group into normal or low CD4 expression at the first blood draw visit. Red dots indicate the 5 participants with low CD4 gMFI in visit 1. Data were analyzed using a 1-way ANOVA with Tukey’s post-test (*\*P < 0.05, **P < 0.01*). **(C)** Percentages of CD69^+^CD134^+^ activated CD3^+^CD8^−^ T cells induced by SEB in PBMCs from two sequential blood draws (1, 2). Paired t-tests were conducted between blood draws 1 and 2. **(D)** Percentages of CD69^+^CD134^+^ activated CD3^+^CD8^−^ T cells correlated with pre-stimulation CD4 expression (gMFI) for all blood draws. Correlation and significance were assessed with a Pearson’s test.

### 3.4 CD4 Downregulation Correlates with Elevated Plasma Cytokines but not Deficient Anti-SARS-CoV-2 Antibody Responses

Next, we investigated whether decreased T cell CD4 expression correlated with specific plasma factors using a multiplexed analysis of 52 mediators. Pearson’s tests revealed that the levels of the following cytokines were inversely correlated with CD4 expression: IL-1RA (r = –0.52, *P*=0.011), IL-7 (r = –0.52, *P*=0.011), and VEGF (r = –0.50, *P*=0.015) **(Fig. 3A-C)**. These findings show that increased concentrations of specific inflammatory mediators are associated with CD4 downregulation and T cell dysfunction observed in individuals with prolonged symptoms. In contrast, Anti-SARS-CoV-2 S1 IgG antibody titers measured in participant plasma samples (n=20) showed no significant correlation with CD4 expression levels (Pearson’s test r = –0.09, *P*=0.680) **(Fig. 3D)**. These data show that the CD4^+^ T cell defect is independent of the humoral response against the virus at the time points assayed.

**Figure 3.**
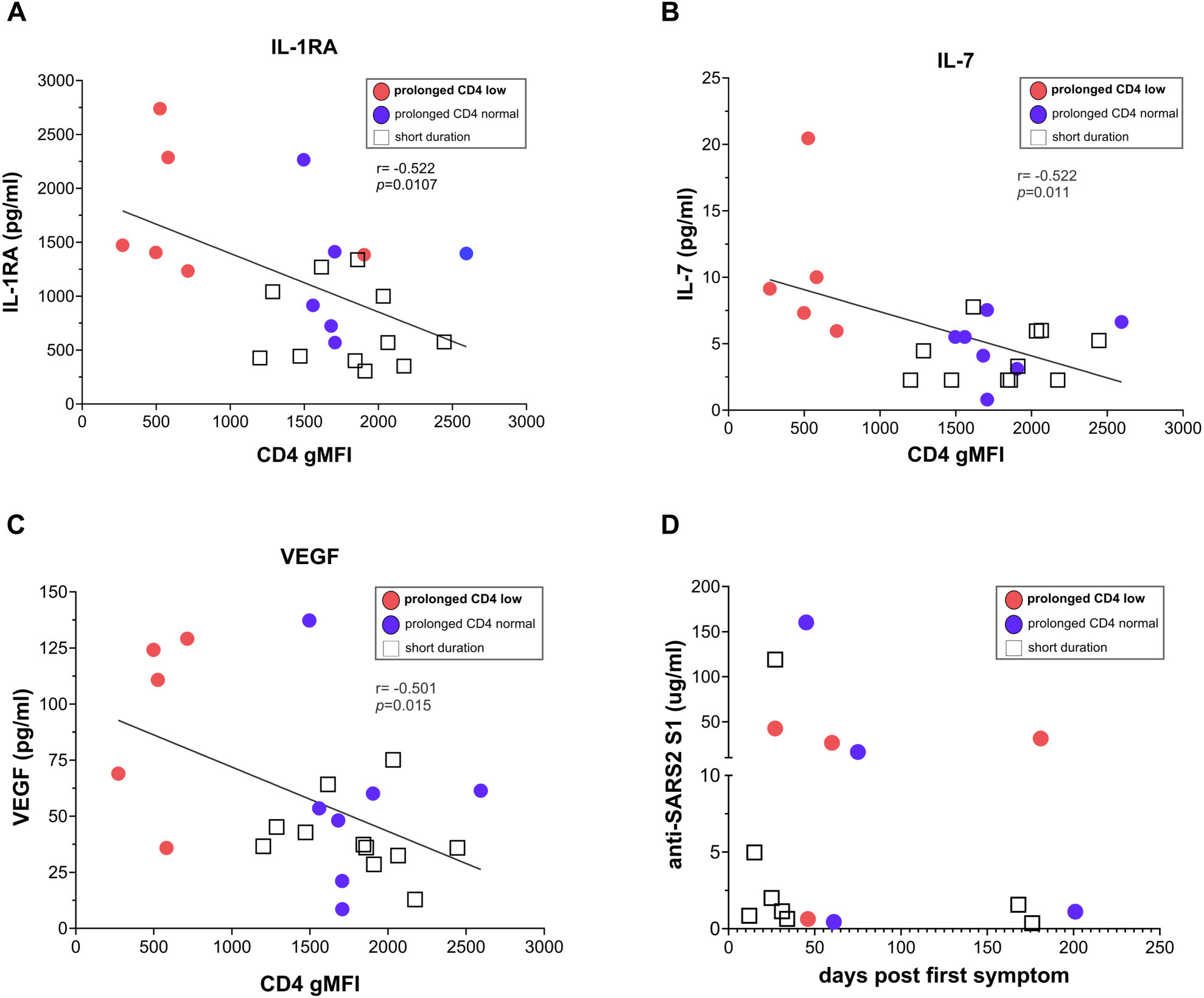
Elevated plasma levels of select cytokines, but not anti-SARS2 S1 antibodies, correlated with reduced T cell CD4 expression. **(A-C)** Plasma cytokines measured for each blood draw in individuals with short or prolonged duration of symptoms using multiplex Luminex assays. Cytokine levels (pg/ml) correlated inversely with CD4 gMFI on CD3^+^CD8^-^ T cells. Significance was assessed with a 2-tailed Pearson’s test. **(D)** Levels of antibody specific for SARS S1 protein measured by ELISA visualized against days after first symptoms reported. Red dots represent data from individuals in the prolonged symptom duration group with low CD4 expression values in visit 1 (n=5). Blue dots represent data from individuals in the prolonged symptom duration group with normal CD4 expression values (n=7). White squares represent data from individuals in the short symptom duration group (n=11).

### 3.5 Transcriptional Profiling Reveals Global Activation Defect in T Cells with low CD4 Expression

To gain deeper insight into the observed functional defects, we performed RNA sequencing of untreated and SEB-stimulated CD3^+^CD8^−^ T cells isolated from the blood of five participants in the prolonged symptom group and from five healthy controls. From the five individuals who initially showed low CD4 expression, we sorted CD3^+^CD8^−^ T cells from one sample when CD4 expression was low and from a second sample when CD4 expression had recovered to normal levels. We labeled these two longitudinal samples as CD4^low^ and CD4^rec^. At a single time point, we also sorted CD3^+^CD8^−^ T cells from healthy controls (HC) matched according to age, race, and sex **(Table 2)**. We conducted bulk RNA sequencing on the sorted cells to identify differentially expressed (DE) genes, defined as those with log_2_FC >1.0 and an FDR-adjusted *P-*value <0.05 **(Fig. S3)**. Strikingly, the CD4^low^ group exhibited 56% fewer DE genes following SEB stimulation than the HC group (1,674 vs. 3,821), and similarly, 57% fewer DE genes than the CD4^rec^ group (1,674 vs. 3,931) (**Fig. 4A; Table S2, S3, S4)**. The CD4^low^ group demonstrated 67% fewer upregulated DE genes than the healthy control group (563 vs. 1,709) and 72% fewer than the CD4^rec^ group (563 vs. 2,034) **(Fig. 4B)**. The CD4^low^ group expressed 47% fewer downregulated genes than the HC group (1,111 vs. 2,112) and 41% fewer downregulated genes than the CD4^rec^ group (1,111 vs. 1,897) **(Fig. 4C)**. These differences indicate a severely blunted transcriptional response to TCR stimulation in participant T cells with low CD4 expression.

**Figure 4.**
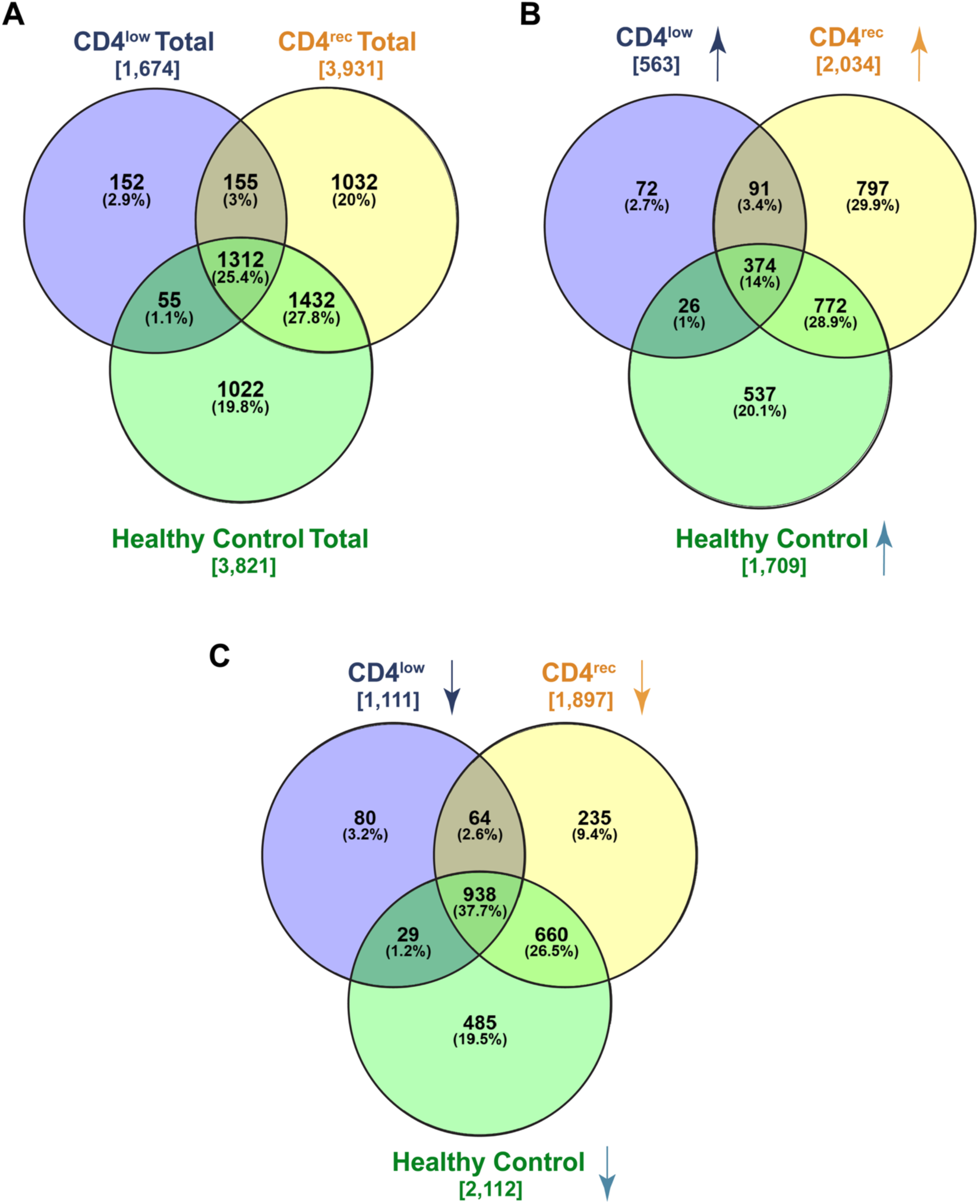
Samples from the CD4^low^ group show >50% fewer differentially expressed (DE) genes after SEB stimulation compared to samples from the CD4^rec^ or Healthy Control groups. Venn diagrams show significant DE genes shared between response groups **(A)** All DE genes **(B)** up-regulated DE genes and **(C)** down-regulated DE genes. DE genes were defined as those with log_2_FC >1.0 and an FDR-adjusted *P-*value <0.05.

**Table 2.** RNA Sequencing Study Participant Demographics. Demographics of healthy controls (n=5) that were age, race, and sex-matched to study participants (n=5) for our RNA Sequencing assays. “n*s”, not significant*.

| Parameters | Healthy Controls (n=5) | Matched Patients (n=5) | P-value |
| --- | --- | --- | --- |
| Age (mean± SD) | 47±16.1 | 46.6±18.2 | ns<sup>a</sup> |
| Race (% White) | 4/5 (80%) | 4/5 (80%) | ns<sup>b</sup> |
| Gender |  |  |  |
| *Female* | 4/5 (80%) | 4/5(80%) | ns<sup>b</sup> |
| *Males* | 1/5 (20%) | 1/5 (20%) |  |

<sup>a</sup> unpaired t-test, <sup>b</sup> Fisher's exact test

### 3.6 Upstream Regulator Analysis Reveals Interferon Signature and Deficient TCR Signaling in CD3^+^CD8^−^ T cells with Low CD4 Expression

To identify the genes and biological pathways of greatest interest, we compared the differentially expressed transcriptome of the CD4^low^ vs. the CD4^rec^ group of unstimulated and SEB-stimulated CD3^+^CD8^−^ T cells **(Fig. 5A, B; Table S5, S6)**. Compared to CD4^rec^ cells, unstimulated CD4^low^ cells showed increased expression of IFN-stimulated genes (*IFI44L, OAS1, IFIT3, and CXCL10*) and the transcription factor *EGR1,* which is reported to be elevated in polarized T_H_2 cells **(Fig. 5A)** [33]. We also noted decreased expression of *SKIL* and *KLF10*, which are involved in regulation of TGFβ signaling in T cells [34–36]. In SEB-stimulated cells, we noted increased expression of *SMAD3* and *VIPR2*, associated with TGFβ and regulatory T cell activity, and of *CD52* and *NUAK2*, associated with decreased T cell activation [37–42] **(Fig. 5B)**. We also noted the presence of several DE genes characteristic of dendritic cells (*CLEC10A* and *CD1C*, Tables S3 and S5) and B lymphocytes (*IGLV8-61* and *IGLV3-13*, Tables S5 and S6), which were possibly present due to doublet contamination among the sorted cells [43].

**Figure 5.**
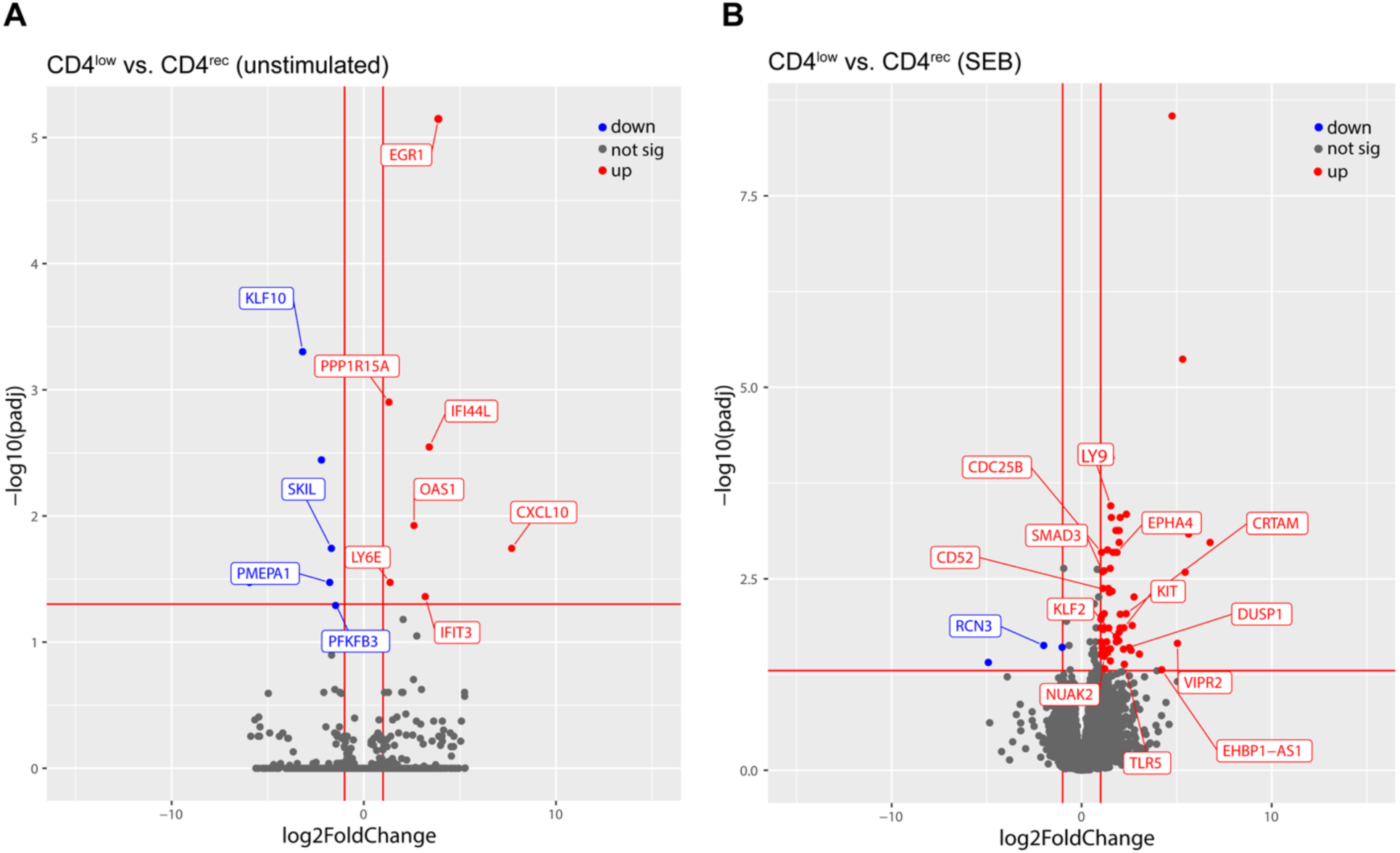
Volcano plots showing selected differential gene (DE) expression between CD4^low^ and CD4^rec^ groups. Shown are plots of log_2_fold change vs. -log_10_(padj). **(A)** DE genes in unstimulated T cells. **(B)** DE genes after SEB stimulation. The up-regulated and down-regulated DE genes are depicted by colors of red and blue, respectively. The non-DE genes are indicated by the gray color. “ns,” not significant.

We next performed Ingenuity Pathway Analysis (IPA) to identify the canonical pathways and predicted upstream regulators **(Fig. 6)** responsible for the observed differentially expressed genes. The top canonical pathway enriched in unstimulated CD4^low^ T cells was the “Hypercytokinemia/hyperchemokinemia in the pathogenesis of influenza” pathway containing multiple IFN-stimulated genes, suggesting prior exposure to an IFN-rich environment in CD4^low^ individuals relative to the CD4^rec^ individuals **(Fig. 6A; Table S7)**. The predicted upstream drivers that could activate this pathway include poly I:C RNA, IFN-α, IFN-β, IFN-γ, TNF, IRF1, IRF3, and STING1. These drivers either mimic viral RNA or respond early to viral RNA [44, 45]. The predicted regulators that would inhibit this pathway include SOCS1, which attenuates IFN signaling pathways **(Fig. 6C; Table S11)** [46].

**Figure 6.**
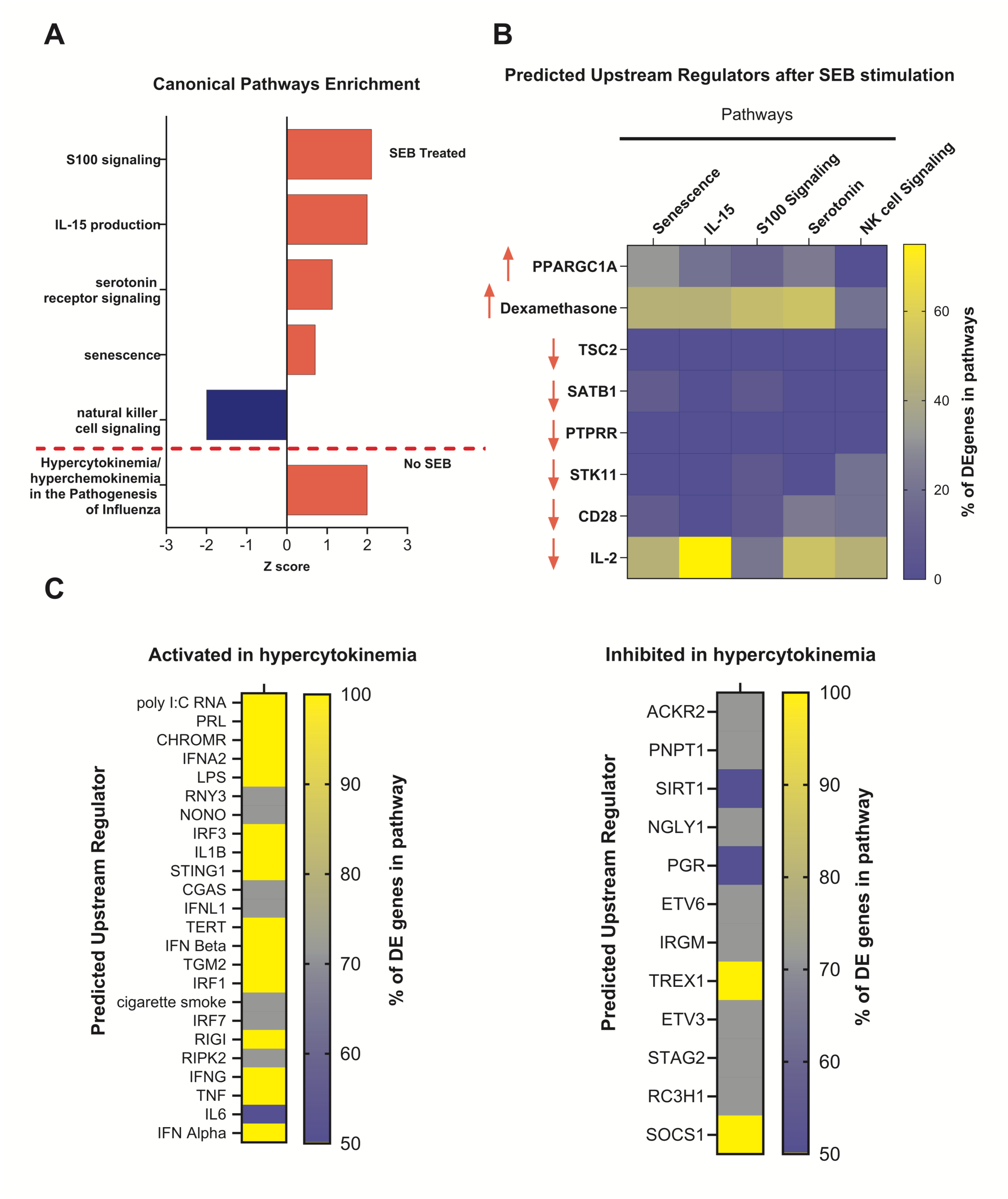
Upstream regulator analyses reveal that unstimulated CD4^low^ T cells were exposed to interferons and that stimulated CD4^low^ T cells show deficient IL-2 and TCR-associated signaling. DE genes in CD3^+^CD8^−^ T cells from CD4^low^ vs. CD4^rec^ groups (see Fig. 5) were subjected to Ingenuity Pathway Analyses. (A) Canonical pathways increased in the unstimulated and SEB-stimulated groups are shown in red. A pathway decreased in the SEB-treated group is shown in blue. (B) SEB-activated T cells: Percentage of DE genes in the indicated pathways (top) that are regulated by the predicted upstream regulator shown (left). Arrows indicate predicted reduced (↓) or increased (↑) influence of the upstream driver in the CD4^low^ group relative to the CD4^rec^ group based on a z-score greater than +/-2. (C) Unstimulated T cells: for the hypercytokinemia and hyperchemokinemia pathways, upstream regulators with a positive z-score greater >2 indicating activation and with a negative z-score < -2 indicating inhibition are shown.

SEB-stimulated CD4^low^ T cells showed increased expression of genes in the pathways “S100 signaling”, “IL-15 production”, “serotonin receptor signaling”, and “senescence” **(Fig. 6A; Table S8)**. The predicted upstream regulators activating these pathways include *PPARGC1A*, a transcriptional coactivator that regulates genes involved in energy metabolism and is associated with senescence [47] and PGC1α, the product of *PPARGC1A*, which forms a complex with GATA3, ATR, and NRF2 to enhance mitochondrial biogenesis, maintaining the viability of CD4^+^ T cells exposed to high reactive oxygen species that can induce DNA damage [48]. Dexamethasone was identified as an upstream regulator predicted to drive the observed gene expression profile, suggesting that T cells within CD4^low^ individuals harbor an immunosuppressed phenotype similar to that imposed by dexamethasone **(Fig. 6B; Table S9, S10).**

## 4. Discussion

Non-hospitalized individuals with prolonged COVID-19 symptoms may exhibit profound alterations in immune function [49]. Herein, we report that a subset of individuals with a prolonged period of symptoms after SARS-CoV-2 infection harbored CD4^+^ T cells with reduced CD4 expression and functional deficits compared to individuals with a short period of symptoms. CD4^low^ T cells exhibited a marked deficit in TCR stimulation via SEB, which was indicated by a failure to upregulate the activation markers CD69 and CD134 compared to CD4^rec^ T cells obtained from longitudinal samples of the same donors or T cells from healthy controls. This TCR signaling hypo-response was reflected in the transcriptome with or without SEB stimulation, with unstimulated CD4^low^ T cells showing evidence of exposure to type I and II IFN and SEB-stimulated CD4^low^ T cells showing evidence of senescence and decreased activation via CD28 and IL-2.

The observed CD4 levels on CD3^+^CD8^−^ T cells showed strong inverse correlations with specific plasma cytokine levels, particularly IL-1RA, IL-7, and VEGF, but notably showed no correlation with anti-SARS-CoV-2 antibody levels. This pattern suggests that persistent immune dysregulation may be driven by ongoing inflammatory processes rather than a humoral immune response deficit. Elevated levels of IL-7, which promote T cell survival and proliferation, potentially represent a compensatory mechanism for the maintenance of T cell homeostasis and function [50, 51]. Consistent with this, SARS-CoV-2-specific CD4^+^ T cells in convalescent individuals have been reported to proliferate homeostatically and persist long term if they expressed detectable levels of IL-7R [15]. Notably, VEGF is associated with severe and long COVID-19 manifestations [30, 52].

Transcriptional profiling of CD3^+^CD8^−^ T cells with low CD4 expression revealed a global activation defect, with a substantially reduced transcriptional response to TCR stimulation compared to cells from healthy control subjects, providing molecular evidence for the observed functional impairments. Upstream regulator analysis identified two key features underlying T cell abnormalities in individuals with prolonged COVID-19 symptoms and low CD4 levels: an IFN signature and deficient TCR signaling. The data showed an increased expression of genes associated with IFN exposure, suggesting prior exposure to an interferon-rich environment. A similar “type I interferon response” module has been reported in blood T cells present in acute samples of participants who subsequently developed long COVID-19 compared to participants who recovered quickly [5]. This interferon signature aligns with previous studies implicating type I IFN in severe COVID-19 pathogenesis and suggests that these effects persist even after viral clearance [53].

Our analysis also predicted inhibition of key TCR signaling components, providing molecular evidence for deficient T cell activation pathways. SEB-stimulated CD4^low^ T cells harbored gene expression profiles consistent with deficient IL-2- and TCR-associated signaling. This profile may arise because the CD4-TCR interactions on activated T cells play a critical role in amplifying the strength of TCR signaling. CD4 stabilizes the interaction of the TCR with MHCII on antigen presenting cells [27]. The kinase LCK, associated with CD4, phosphorylates the CD3 zeta chain at residues 21 and 23. The phosphorylated zeta chain residues recruit ZAP70, which is also phosphorylated by LCK. These events promote the signal transduction cascade downstream of the TCR that occurs during T cell activation. However, CD4 is not required for activation of T cells by SEB. The superantigen SEB can bypass the CD4-LCK-dependent TCR signaling pathway and activate a noncanonical pathway involving the G protein Gαq or Gα11 and the phospholipase Cβ protein PLCβ2 in human T cells [28, 29]. Signaling via this alternate pathway leads to IL-2 production via a PLCβ and PKC mediated cascade that results in ERK signaling, Ca^2+^ influx, and nuclear translocation of NF-AT and NF-κB. Notably, we found that SEB stimulation of CD4^rec^ T cells led to a 2-fold increase in expression of *PLCB2* (encoding PLCβ2), and this increase was not observed upon SEB stimulation of CD4^low^ T cells. These data suggest that the CD4^low^ T cells harbor functional deficits that would impact both CD4-LCK dependent signaling in response to MHCII-presented peptide antigens as well as LCK-independent TCR signaling triggered by the superantigen SEB.

Reduced CD4 on T cells has been associated with chronic *in vivo* or *in vitro* stimulation, T cell exhaustion, and IFN-driven autoimmune diseases such as systemic lupus erythematosus [26]. The CD4^low^ T cells also harbored expression of genes indicating exposure to IFNs, which has been shown to enhance and maintain PD-1 expression after TCR engagement [27]. However, unstimulated CD4^low^ T cells did not show elevated surface PD-1 or CD25 expression, and they did not show differential expression of canonical T cell exhaustion genes, including *TIM3, LAG3, PD1, TIGIT*, suggesting that the CD4^low^ T cells are not exhausted by these criteria (*data not shown*). Unstimulated CD4^low^ T cells did show reduced expression of *PFKFB3*, encoding phosphofructokinase, known to be involved in glycolysis. Studies of CD4^+^ T cells in individuals with rheumatoid arthritis showed that insufficient PFKFB3 impaired ATP generation and promoted senescence [54]. SEB-stimulated CD4^low^ T cells showed increased expression of *CD52* and *NUAK2*, both encoding proteins that interfere with or suppress T cell activation [39, 41, 42].

T cells bearing reduced CD4 are hyporesponsive to stimulation, which may reflect chronic inflammation but also may direct alternate fates [55]. For example, a study of polarized T_H_1 and T_H_2 cells bearing the same TCR showed that T_H_2 cells expressed lower levels of CD4, which is linked with decreased TCR-associated protein tyrosine phosphorylation with the absence of ZAP70 phosphorylation and partial phosphorylation of TCR zeta residues [56]. Enforced CD4 expression in T_H_2 cells restored the zeta chain and ZAP70 phosphorylation to levels found in T_H_1 cells, suggesting that the CD4-LCK complex regulates the extent of TCR-mediated activation.

Interestingly, we observed that relative to CD4^rec^ T cells, unstimulated CD4^low^ T cells expressed high levels of *EGR1*, which is preferentially expressed in T_H_2 cells [33]. Consistent with our findings, a recent study showed that compared to individuals who had recovered shortly after acute infection, the blood of individuals with long COVID-19 was enriched for T cells that harbored higher amounts of intracellular IL-4 at 3 and 12-months post-infection. These data suggest that development of long COVID-19 is associated with a predisposition to T_H_2 immunity [5].

In addition, a pattern of gene expression in CD4^low^ T cells compared to CD4^rec^ T cells suggested differentiation along the T regulatory cell pathway at the expense of T_H_1 pathways. Unstimulated CD4^low^ T cells showed decreased expression of *SKIL* (known as *SnoN* in mice), a negative regulator of antiproliferative TGFβ signaling [34]. Murine T cell deficiency of *Skil/SnoN* leads to increased TGFβ signaling, regulatory T cell differentiation, and decreased T cell proliferation in response to activating stimuli [35]. SEB-stimulated CD4^low^ T cells showed increased expression of *SMAD3*, which is essential for TGFβ-mediated induction of *FOXP3* [37]. CD4^low^ T cells also showed increased expression of *VIPR2*, and it has been reported that VIPR2 agonists promote regulatory T cell activity [38]. These data are consistent with the prior finding that severe infection with poor outcomes is associated with fewer SARS-CoV-2-specific CD4^+^ T_H_1 and T_FH_ cells and a higher number of T regulatory cells [16].

The identification of persistent CD4^+^ T cell dysfunction raises important clinical issues that warrant further investigation. Our study demonstrates that such dysfunction occurs in non-hospitalized individuals with prolonged symptoms, indicating that even a moderate COVID-19 infection course can lead to significant immune alterations. Understanding lymphocyte phenotypes over the timeline of recovery is crucial, as our study shows individual variation in recovery patterns, with some individuals showing delayed restoration of normal T cell function extending well beyond the acute infection period. The potential impact of these prolonged defects on susceptibility to other infections, the overall CD4^+^ T cell memory pool, and the efficacy of vaccinations in affected individuals require further exploration to reveal the clinical implications of these findings.

## CRediT reporting

Jadith Ziegler – Formal analysis, Visualization, Writing – original draft, Writing – review & editing

Christina Lawrence – Investigation, Writing – review & editing

Sean Turner – Investigation, Writing – review & editing

Nathan Pezant – Formal analysis, Visualization, Writing – review & editing

Nancy Redinger – Resources, Writing – review & editing

Carla Guthridge – Investigation, Writing – review & editing

Kenneth Smith – Investigation, Writing – review & editing

Jingrong Chen – Investigation, Writing – review & editing

Joel M. Guthridge – Resources, Writing – review & editing

Judith A. James – Funding acquisition, Resources, Writing – review & editing

Susannah Rankin – Resources, Writing – review & editing

Linda F. Thompson – Conceptualization, Funding acquisition, Writing – review and editing

Darise Farris – Conceptualization, Funding acquisition, Project Administration, Formal analysis, Supervision, Writing – review and editing

Susan Kovats – Conceptualization, Funding acquisition, Project Administration, Formal analysis, Supervision, Visualization, Writing – original draft, Writing – review and editing

## Data availability

The RNA sequencing data were deposited in Mendeley: Kovats, Susan; Farris, A. Darise (2025), “CD4 T cell covid-19 Ziegler et al”, Mendeley Data, V1, reserved doi: 10.17632/8p2pzwbgw5.1

## Acknowledgements

We thank the OMRF Flow Cytometry Core Facility for expert cell sorting and the OMRF Clinical Genomics Center for RNA sequencing.

## Supplemental Figures and Tables

**Table S1:** Additional study participant information

**Table S2:** Differentially expressed genes in healthy control CD3^+^CD8^-^ T cells (SEB stimulated/unstimulated)

**Table S3:** Differentially expressed genes in CD4^rec^ participants’ CD3^+^CD8^-^ T cells (SEB stimulated/unstimulated)

**Table S4:** Differentially expressed genes in CD4^low^ participants’ CD3^+^CD8^-^ T cells (SEB stimulated/unstimulated)

**Table S5:** Differentially expressed genes in CD4^low^ group relative to CD4^rec^ group without SEB stimulation

**Table S6:** Differentially expressed genes in CD4^low^ group relative to CD4^rec^ group after SEB stimulation

**Table S7:** Significant canonical pathways and their involved genes in CD4^low^ group relative to CD4^rec^ group without SEB stimulation

**Table S8:** Significant canonical pathways and their involved genes in CD4^low^ group relative to CD4^rec^ group after SEB stimulation

**Table S9:** Differentially expressed upstream regulators predicted by IPA in CD4^low^ group relative to CD4^rec^ group without SEB stimulation

**Table S10:** Differentially expressed upstream regulators predicted by IPA in CD4^low^ group relative to CD4^rec^ group after SEB stimulation

**Table S11:** Differentially expressed upstream regulators of hypercytokinemia predicted by IPA in CD4^low^ group relative to CD4^rec^ group without SEB stimulation

**Supplemental Figure 1.**
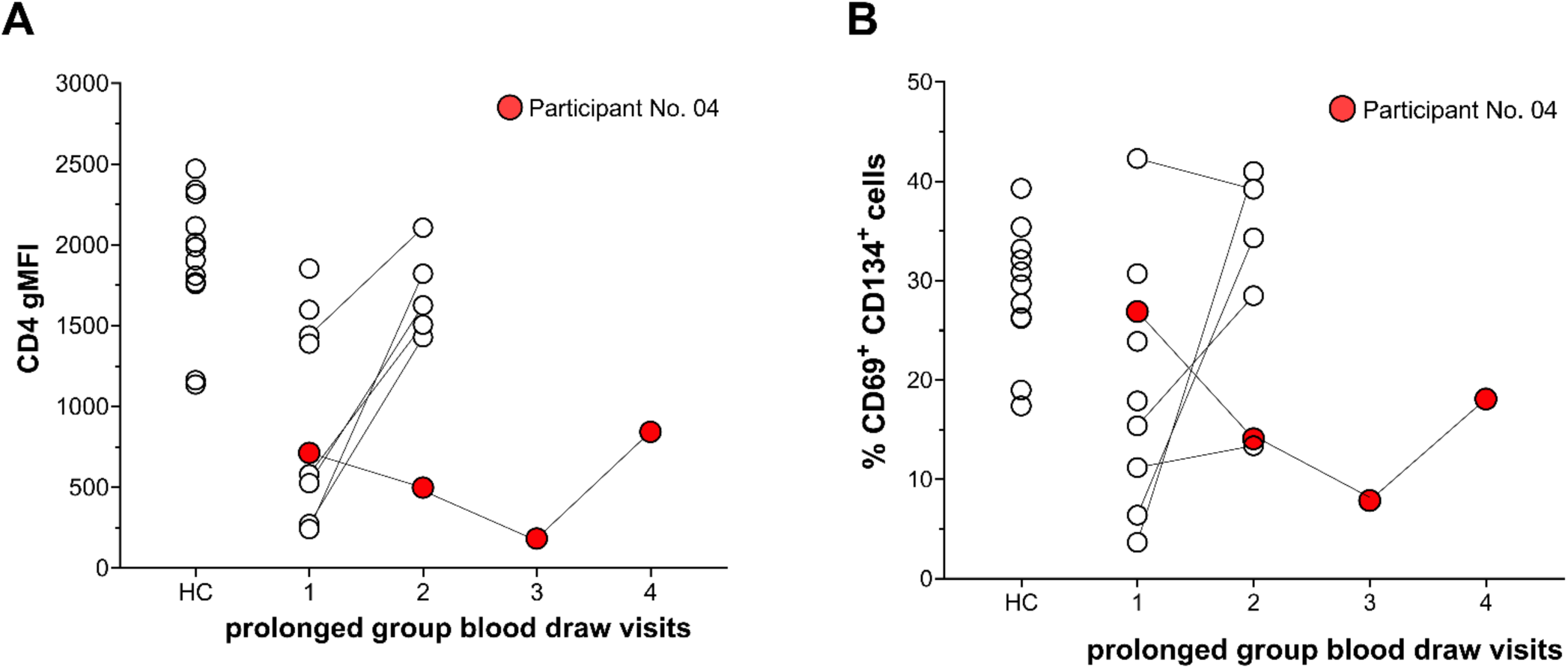
**(A)** CD4 gMFI values and **(B)** % CD69^+^ CD134^+^ CD3^+^CD8^−^ T cells after SEB stimulation including data from all blood draws from participant 04 (in red).

**Supplemental Figure 2.**
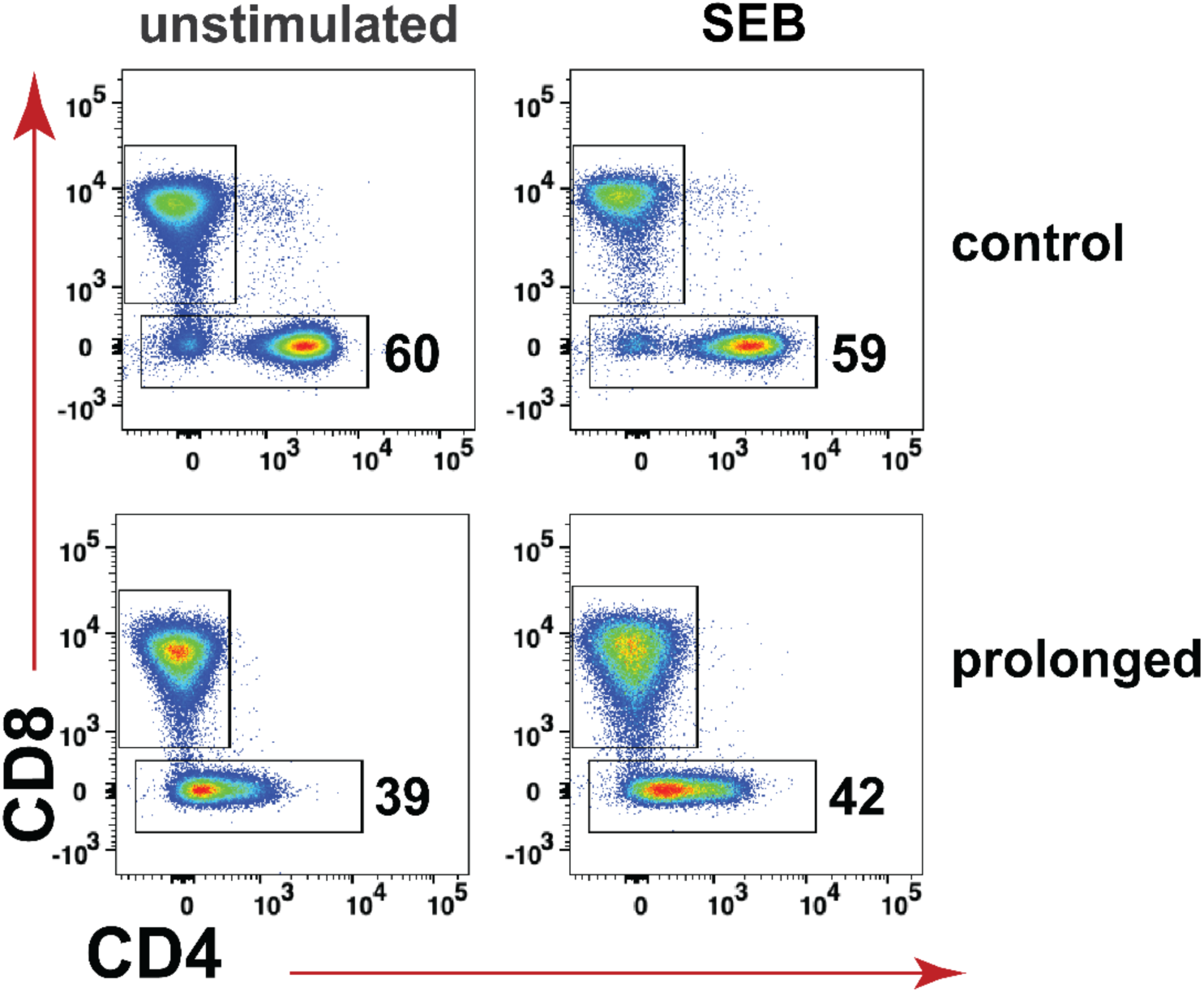
T cell sorting gating strategy used for RNA Sequencing.

**Supplemental Figure 3:**
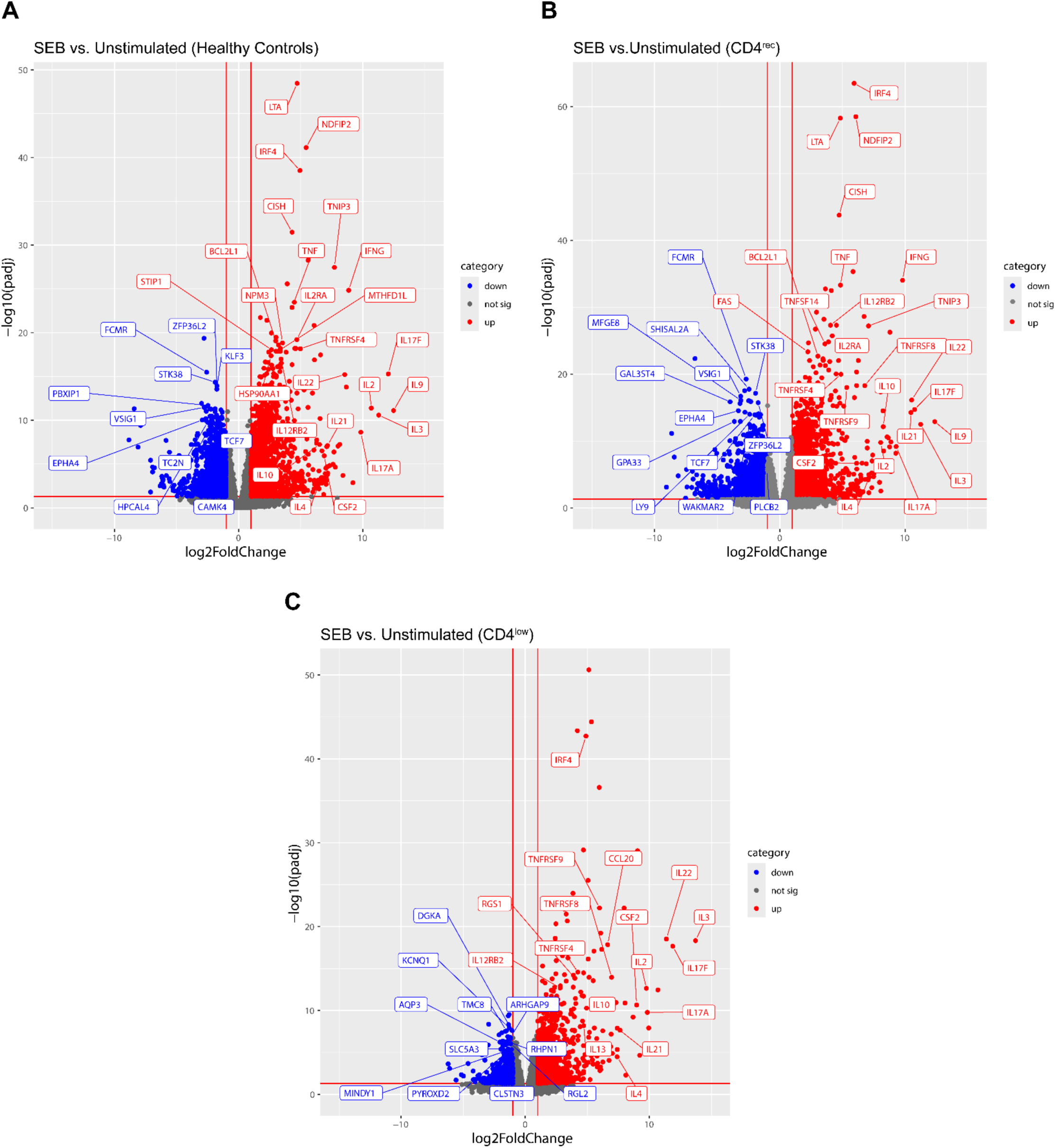
Volcano plots of top differently expressed genes comparing unstimulated versus SEB stimulated within sample groups **(A)** healthy control, **(B)** CD4^rec^, and **(C)** CD4^low^ groups.

